# *In Vivo* Subcellular Mass Spectrometry Enables Proteo-Metabolomic Single-cell Systems Biology in a Chordate Embryo Developing to a Normally Behaving Tadpole (*X. laevis*)

**DOI:** 10.1101/2021.01.19.426900

**Authors:** Camille Lombard-Banek, Jie Li, Erika P. Portero, Rosemary M. Onjiko, Chase D. Singer, David Plotnick, Reem Q. Al Shabeeb, Peter Nemes

## Abstract

We present the first example of *in vivo* high-resolution mass spectrometry (HRMS) for subcellular molecular systems biology of proteins and metabolites. With light microscopy, we identified the left-dorsal and left-ventral animal cells in cleavage-stage non-sentient *Xenopus laevis* embryos. Using precision-translated fabricated microcapillaries, the subcellular content of each cell was double-probed, each time collecting <5% of cell volume (~10 nL) swiftly (<5 s/event). The proteins and metabolites were analyzed by custom-built ultrasensitive capillary electrophoresis electrospray ionization employing Orbitrap and time-of-flight HRMS. Label-free detection of ~150 metabolites (57 identified) and 738 proteins found proteo-metabolomic networks with differential quantitative activities between the cell types. Spatially and temporally scalable sampling the technology preserved the integrity of the analyzed cells, the neighboring cells, and the embryo. 95% of the analyzed embryos developed into sentient tadpoles that were indistinguishable from their wild-type siblings based on anatomy and visual function in a background color preference assay.

---

Unbiased measurement of transcripts, proteins, and metabolites in the live cell is the holy grail of molecular systems cell biology. Even today, after the invention of single-cell transcriptomics^1–4^, there exists no single technology capable of the unbiased characterization of both proteins and metabolites in the same single cell *in vivo* to enable the study of live organisms. While tools of molecular biology and high/super-resolution optical microscopy empowered systems biology for live organisms, they only work for a limited number of gene products at a time that also have to be known a priori, thus limiting research scope to targeted studies, typically building on prior knowledge. Singlecell high-resolution mass spectrometry (HRMS) emerged as a powerful alternative for targeted as well as untargeted (discovery) studies (reviewed in refs. ^1–2, 4–7^), yielding data to derive new hypotheses for biological investigations. Specialized technologies in cell handling, sample processing^8–13^, high-efficiency separations^14–16^, and ion generation equipped time-of-flight and orbitrap HRMS instruments with exquisite sensitivity. For example, using nanoHPLC, NanoPOTS^13^ enabled the detection of 650+ proteins in single HeLa cells, which was recently increased to 850+ proteins on a new-generation mass spectrometer^16^. ScoPE-MS also used nanoHPLC to quantify ~750 proteins from a mammalian cell.^12^ The technology was recently advanced to the quantification of ~1,000 proteins/cell.^12, 17^ We and others developed ultrasensitive capillary electrophoresis (CE) electrospray ionization (ESI) platforms to detect transcripts, hundreds of proteins, or ~50 metabolites in single cells/neurons dissected from *Aplysia californica*^14, 18–20^, *Xenopus laevis* embryos^8, 21–22^, or the mouse^14^. Using capillary microsampling, we extended these measurements to the direct analysis of metabolites^23–24^ or proteins^11^ in single cells in early developing *X. laevis* and zebrafish embryos. MALDI-TOF followed by nanoLC HRMS recently enabled the characterization of lipids, peptides, and proteins in dissociated single cells of single dorsal root ganglion cells from the mouse.^25^ Although these and other technologies usher an exciting new area of multi-omic analyses in single cells,^26–27^ these HRMS tools required isolation or sorting of the cells, preventing studies *in vivo*. We here report the first example of *in vivo* single-cell HRMS that enables dual proteo-metabolomics of spatiotemporally identified single cells in a live embryo freely developing to a normally behaving tadpole post analysis.

We developed the bioanalytical technology and demonstrated its use in molecular systems cell biology with compatibility for cell-, neurodevelopmental, and behavioral biology. **Figure 1** presents our analytical and biological tracks, essentially connecting the physical and life sciences. The former is designed to yield quantitative single-cell multi-omics, with an option for cell identification in space and time, to enable molecular network analyses based on detected proteins and metabolites. The latter is purposed to assess the biological phenotype via morphological, survival, and behavioral assays, thus supporting hypothesis-driven studies in biology and health research. By design, the integration of these tracks generates multi-dimensional metadata to open a window into molecular systems biology and help develop new hypotheses and knowledge.

**Figure 1.**
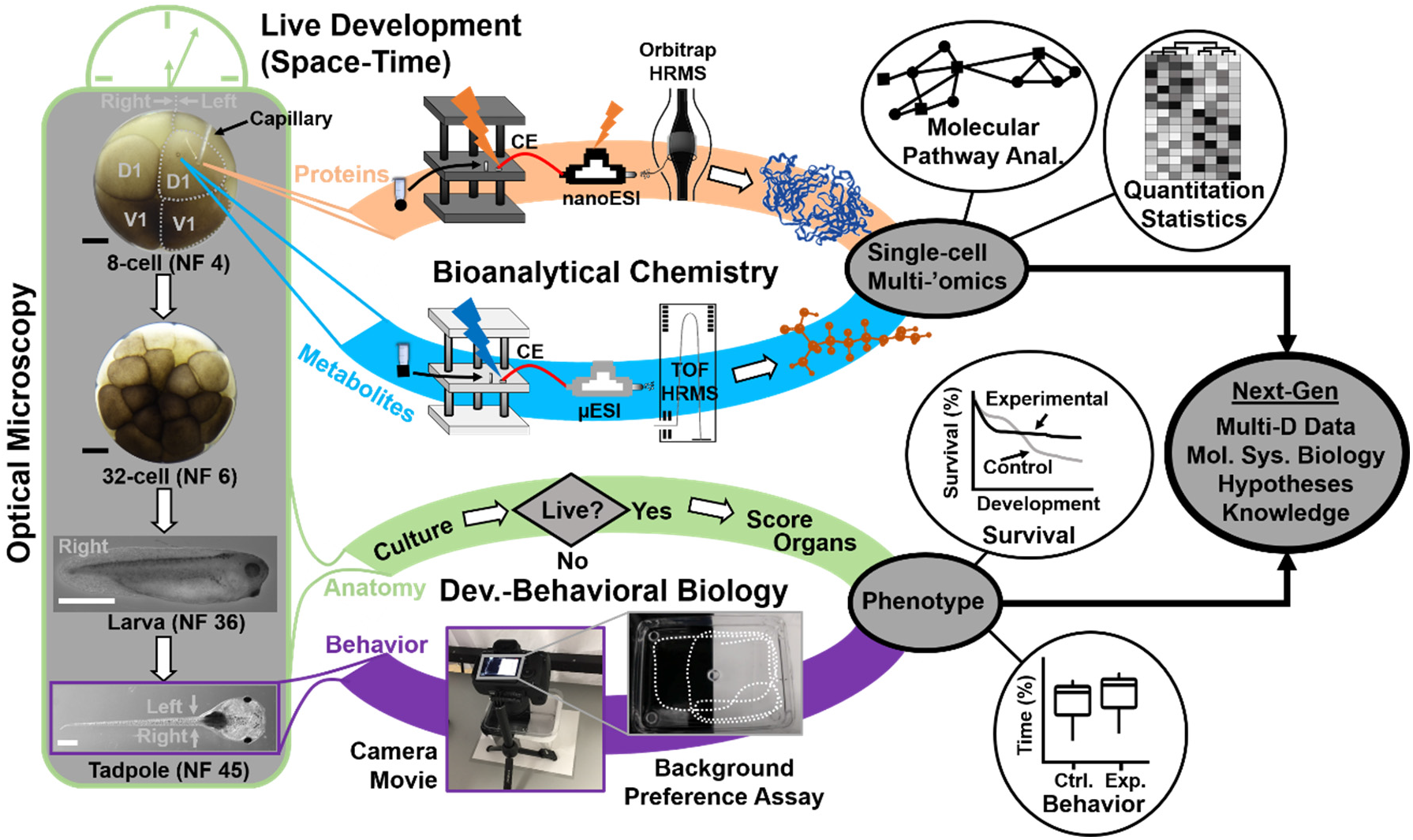
Interdisciplinary strategy enabling *in vivo* subcellular proteo-metabolomic systems biology with compatibility for cell-, neurodevelopmental, and behavioral biology using *Xenopus laevis* as the biological model. The (left) dorsal-animal (D1) and ventral-animal (D1) cells were identified and the content of each cell rapidly microsampled twice. The collected protein and metabolite samples were analyzed using custom-built ultrasensitive capillary electrophoresis (CE) nano/micro-flow electrospray ionization (ESI) high-resolution orbitrap and time-of-flight (TOF) mass spectrometry (HRMS). The tadpoles developing from the embryos were characterized for survival, anatomy, and behavior. Scale bars, 250 μm (black), 1 mm μm (white).

The vertebrate South African clawed frog (*Xenopus laevis*) was an ideal biological model for technology development, validation, and application in this study. Each cell in the cleavage-stage frog embryo is identifiable based on pigmentation and location and has reproducible tissue fates.^28^ **Figure 1** exemplifies the identification of the left dorsal-animal (L-D1) and left ventral-animal (L-V1) cell in a live 8-cell embryo, which respectively form the neural and epidermal tissues. *X. laevis* embryos develop externally to the mother and contain considerably large cells (180 nL volume/cell in the 8-cell embryo), permitting direct inspection and access to the cells. However, the cells populate the embryo in a complex three-dimensional morphology and divide rapidly, within ~15– 25 min per cell cycle between the 2- and 32-cell stages.^29^ These characteristics fundamentally challenge existing single-cell HRMS technologies in terms of scalability for spatial, temporal, and *in vivo* operation, all key aspects for studying cell biology in live tissues, organs, and organisms, such as the live *X. laevis* embryo selected here. While dissociation^30^ and manual dissection^8, 21, 31^ permit cell isolation from the embryo, the former requires substantial dexterity and lacks scalability and the latter loses spatial information on cell heterogeneity, which has important biological implications. With operation *ex vivo*, these tools also exclude the possibility of biological studies over critical developmental events, ranging from gastrulation and neurulation to organogenesis and metamorphosis as well as assessment of function and behavior at an organismal level, such as the tadpole as demonstrated in this study (see images in **Fig. 1**). Currently, there exists no single-cell HRMS technology capable of determining the proteo-metabolome of single cells embedded in complex tissues or organisms *in vivo* with scalability.

We proposed double capillary microsampling as a potential solution for analyzing the cells *in vivo*. A microfabricated capillary was connected to a microinjector to deliver negative pressure pulses and the tip of the capillary was inserted into the cell of interest under translation by a precision micromanipulator under guidance by real-time stereomicroscopy. In each of N = 20 stereotypical 8-cell embryos (experimental group), the L-D1 and L-V1 cells were each microsampled twice, both from the same embryo, one for each ‘omic analysis. By performing the single-cell analyses exclusively on one side of the embryo (the left, experimental), tissues and organs arising from the nonsampled cells on the other side (right D1, R-D1 and right V1, R-V1) served as the internal control in each embryo to facilitate analysis of tadpole anatomy (see later). For the control group (Ctrl.), N = 20 stereotypical 8-cell embryos were cultured under identical conditions, without microaspiration. This approach was inspired by *in situ* microprobe sampling that we recently developed for cells in zebrafish and *X. laevis* embryos, albeit functioning only *ex vivo* and exclusively for metabolomics^10, 23–24^ or proteomics^11^, but not both ‘omes at the same time or on the same cell. Technical details are discussed in the **SI** document. We experimentally tested that double microsampling using larger capillaries (e.g., ~80 μm tips) and/or longer aspiration times (scalability) allowed us to aspirate ~100 nL, viz. 50% of the cell volume. These amounts were sufficient for single-cell HRMS; however, the embryos failed to survive the analysis.

To enable *in vivo* cell sampling, we scaled the approach with assistance from survival assays (**Fig. 2**). In a series of experiments (data not shown), the tip size of the microprobe, the volume collected, and the speed of microsampling were tailored to the cells while monitoring the success of cell divisions after sampling under the microscope. Each sampling employed a clean and unused capillary to aid cell and embryonic survival by avoiding accidental contamination. **Figure 2A** tracks the percentage of embryos surviving after ~10 nL, or ~5% of the cell volume, was microaspirated twice rapidly, in less than 30 s/sampling/cell, using 20-μm-tip capillaries from the cells (N = 20). Survival was quantified over 6 key stages of vertebrate development: cleavage (Nieuwkoop-Faber, NF 6), gastrula (NF 10.5, **Fig. 2B**), neurula (NF 13, **Fig. 2B**), early tailbud (NF 22), late tailbud (NF 36 and 41), and tadpole (NF 45, **Fig. 2B**). Compared to 100% of tadpoles successfully developing in the control group (unperturbed wild-type, WT) at all these stages of development, 100% of the embryos developed to the neurula stage and 95% survived to tadpoles in the experimental group. Therefore, capillary microsampling was successfully scaled to preserve viability.

**Figure 2.**
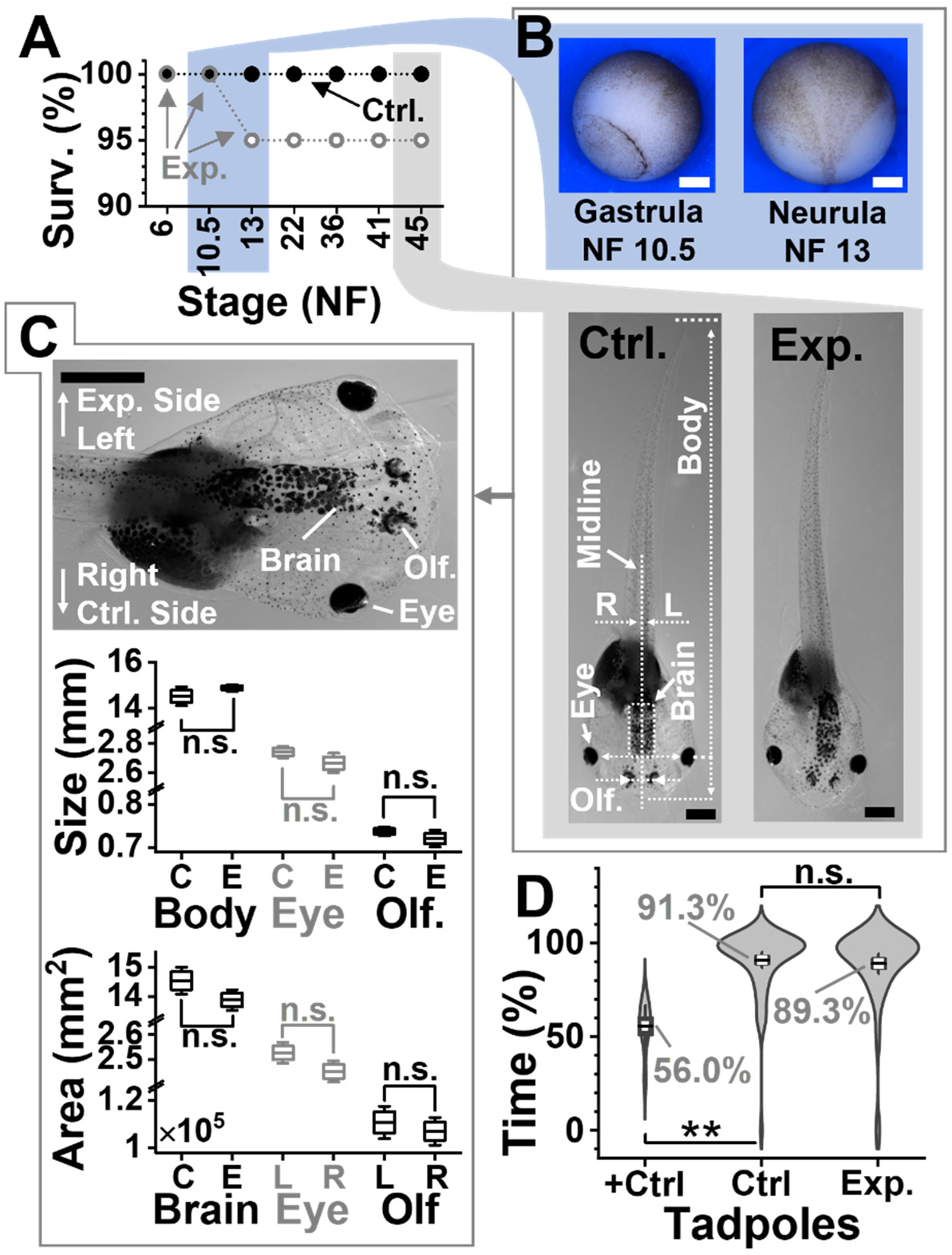
Developmental and behavioral impact of the technology. (**A**) Survival analysis (Surv.) at key developmental stages (NF, Nieuwkoop-Faber) for N = 20 embryos in the control (Ctrl.) group and N = 20 embryos in the experimental. (**B**) Representative (**top panel**) gastrulae and neurulas and (**bottom**) tadpoles. Shown: midline separating left (L)- right (R) axis, body length, brain area, and distance between eyes and olfactory (Olf.) organs. (**C**) Comparison of tadpole anatomy. (**Top**) Close-up image of a tadpole showing the experimental (left) and control (right) side. Left tissues noted. (**Middle**) Analysis of organ size on the left (experimental side) between N = 5 Ctrl. (C) and N = 6 Exp. (E) surviving tadpoles randomly selected. Key: n.s., non-significant, \**p* < 0.05, Mann-Whitney test. (**Bottom**) Analysis of tissue areas for total brain (Ctr. vs. Exp. group) and left-right eye and Olf symmetry for the 6 Exp. tadpoles. Key: n.s., non-significant, Wilcoxon signed rank, paired). (**D**) Comparison of visual function in a background-color preference assay for N = 16 Ctrl. vs. N = 15 Exp. tadpoles. Assay validation via double axotomy of the optic nerves in N = 4 positive Ctrl. tadpoles (+Ctrl.). Tadpole movies in **Videos S1** (Ctrl.) and **S2** (+Ctrl.). Key: n.s., non-significant; \**p* < 0.05, \*\**p* < 0.005 (Mann-Whitney). Scale bars, 250 μm (white) and 1 mm (black). Box-whisker plots: Box, 1×standard error of the mean (SEM), whiskers: 1.5×SEM.

Anatomy was also analyzed for the surviving tadpoles. **Figure 2B** presents typical examples for tadpoles from the control and experimental groups. Their morphologies were characterized in terms of whole-body length as well as the sizes of the eyes, olfactory organs (Olf.), and total brain, which partially derive from the L-D1 and L-V1 cells (**Fig. 2B**). Body symmetry was also analyzed for the eyes and olfactory tissues along the midline separating the left (experimental) side and the right (control) side (see close-up in **Fig. 2C**). As shown in **Figure 2C**, body length (*p* = 0.315, Mann-Whitney) as well as the center-to-center distances between the eyes (*p* = 0.411, Mann-Whitney) and the olfactory tissues (*p* = 0.523, Mann-Whitney) were indistinguishable, as was the total area of the brain (*p* = 0.121, Mann-Whitney). The size (area) of the eyes (*p* = 0.142, Wilcoxon signed rank, paired) and olfactory organs (*p* = 0.142, Wilcoxon signed rank, paired) were also indistinguishable between the experimental and control sides in the tadpoles. Therefore, double subcellular capillary microsampling of the both L-D1 and L-V1 in the cleavage-stage embryo led to no detectable impairment on tadpole development, morphologically.

Physical appearance, however, does not equate to performance; therefore, we further evaluated the animals based on behavior. The visual (also including motor, sensory, and neural processing) function was compared at stages 45–50, when tadpoles display a robust preference for lighter background in a color preference assay^32^ (see setup in **Fig. 1**). Technical details are available in the **SI** document. Indeed, as shown in **Figure 2D**, when presented with a dark vs. light background in a tank, the control (WT) tadpoles resided ~91% of the time over the light background (N = 16). These tadpoles recapitulated this robust behavioral phenotype at this stage,^32^ thus confirming the robustness of the assay in our hands. We further validated the sensitivity of the assay to detect visual impairment using a positive control (+Ctrl). Tadpoles that underwent double axotomy of their optic nerves (N = 4) explored the light background only at 56% of the time, in a significant difference from the control group (*p* = 1.64×10^-5^). Indeed, without vision, tadpoles make random explorations, spending ~50% on either side. These results also agreed with results from an independent investigation^32^, thus confirming that the assay was sufficiently sensitive to detect visual function in our experiments. Swimming ~89% time over the light area, the experimental group (N = 15) presented indistinguishable visual behavior from the control tadpoles (*p* = 0.41, Mann Whitney). Representative behaviors are available for a Ctrl. and +Ctrl. tadpole in **Videos S1 and S2**, respectively. Double microsampling of each of the 2 cells in the same embryo preserved embryonic viability and organismal behavior. The approach opened, for the first time in single-cell HRMS, the possibility of dual ‘omics of single cells in a live organism, which also undergoes dynamic spatial and temporal development to form a behaving animal, as demonstrated here for the *X. laevis* embryo and its tadpole *in vivo* in this work.

The limited amounts of proteins and metabolites from the cells necessitated ultrasensitive detection. From each cell type in the same embryo (N = 4), the double aspirates were independently processed via a proteomic and metabolomic workflow (recall **Fig. 1**). To eliminate potential systematic biases during sample collection, we randomized the location of subcellular sampling and the order of aspirating the protein and metabolite samples. Each embryo, each cell, and each protein and metabolite sample was assigned a unique identifier in our study, although this information was purposefully not used during data analysis to eliminate potential (conscious or unconscious) bias and only revealed to aid the interpretation of results. The bioanalytical workflow entailed the processing of the aspirated samples for microscale bottom-proteomics and metabolomics, separation of the resulting peptide and metabolite samples, efficient ionization, and detection–quantification by HRMS–MS/MS. Technical details are in the **SI** document. The traditional proteomic steps of desalting, alkylation, reduction were eliminated to minimize protein losses (e.g., on vials and pipette tips). The final subcellular samples contained the tryptic digest in 2 μL for bottom-up proteomics and 4 μL for metabolomics. To avoid systematic bias during instrumental measurement, the subcellular ‘omes were blinded for cell type and analyzed in a randomized order. With ~100–10,000-times smaller amounts of material collected from the cells in this study than those typically processed/measured by standard HPLC-HRMS, we turned to our ultrasensitive CE-ESI-HRMS instruments for assistance. These platforms provided ~700 zmol sensitivity for peptides using a tribrid quadrupole-orbitrap-ion trap^8, 11^ and ~60 amol sensitivity for metabolites using a quadrupole time-of-flight high-resolution^21, 23^ mass spectrometer. Technical details are in the **SI** document. To facilitate the instrumental measurements, the metabolic track was limited to cationic electrophoresis in this study, although dual cationic-anionic metabolomics is also possible on these instruments.^33^

Measurement of ~0.5% (v/v) of the protein extract, thus ~0.03% of the cell’s volume, identified 738 proteins (false discovery rate < 1%), listed in **Table S1A**. Similarly, analysis of ~0.25% (v/v) of the metabolite sample (volume), viz. ~0.01% of the cell’s volume, produced ~150 nonredundant metabolic molecular features, 57 of which were identified with a high Level-1^34^ confidence, listed in **Table S1B**. Notably, detection of these ~800 proteins and metabolites was achieved in an untargeted (discovery) manner, requiring no prior knowledge of cellular composition or the use of functional probes; no antibodies, no antisense oligos were necessary for our technology.

These multi-omics data also opened a window into (sub)cellular biochemistry, directly in the live embryo. Statistical enrichment analysis using the Kyoto Encyclopedia of Genes and Genomes^35^ revealed significant coverage of several known pathways (**Fig. 3A**). For example, the arginine-proline pathway, TCA cycle, glycolysis, and pyruvate metabolism were represented with statistical significance, whereas several other amino acid pathways had high pathway impact. Notably, these proteins and metabolites assessed cellular biochemistry in complementary performance. For example, the biosynthesis of aminoacyl-tRNA and valine-leucine-isoleucine was only represented by metabolites, whereas glycolysis/gluconeogenesis was solely enriched for by proteins (**Table S2**). Further, other pathways, such as arginine biosynthesis and arginine-proline metabolism, were represented by both ‘omes (**Table S2**). **Figure 3B** marks the detected proteins and metabolites in the latter. The observed coverage of the network raises a potential for studies targeting particular ‘omic processes. By also including anionic metabolites during CE-ESI-HRMS, we anticipate many other cellular pathways to be detectable by this approach, such as the TCA cycle and energy production as well as drug metabolism. These results illustrate the benefit of measuring more than one ‘ome in single cells toward a holistic understanding of cellular systems biology at the molecular level.

**Figure 3.**
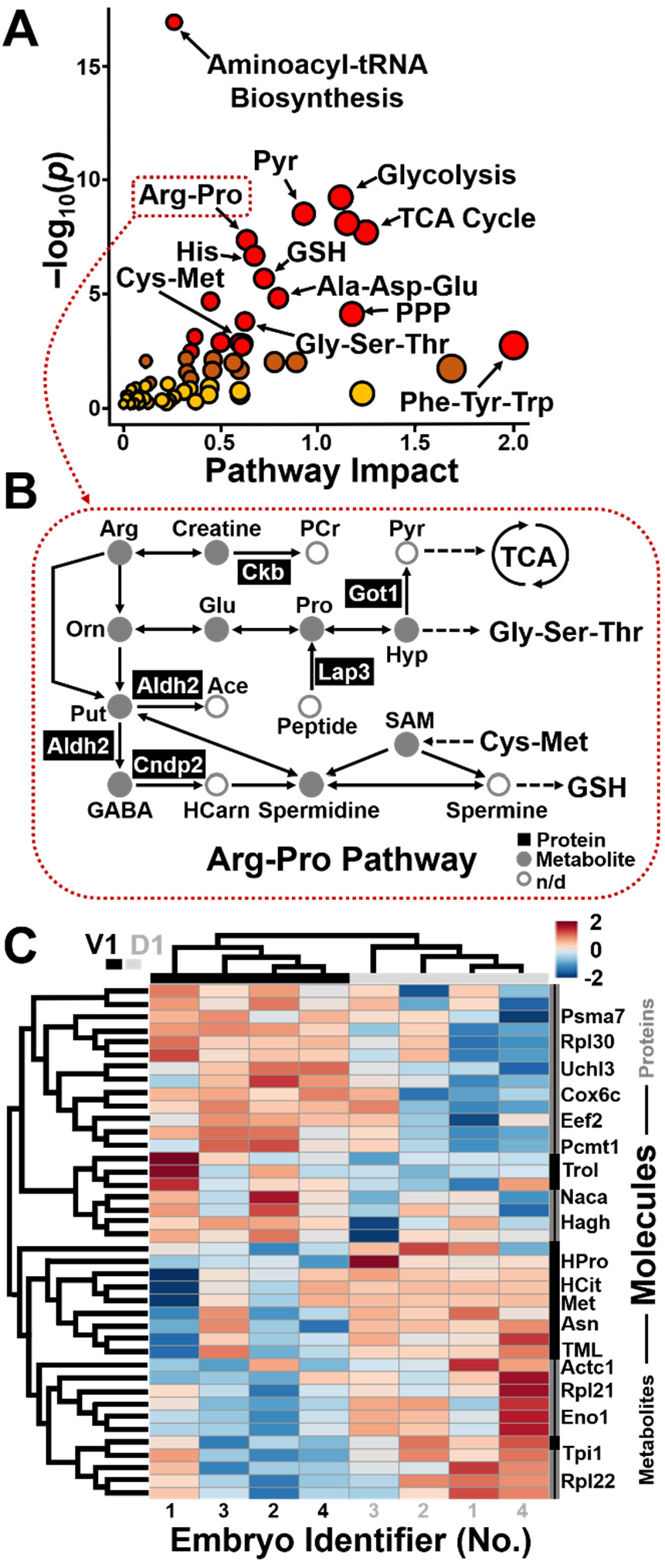
Qualitative and quantitative proteo-metabolomic systems biology on the L-D1 and L-V1 cell in the live embryo based on the *in vivo* single-cell HRMS data. (**A**) KEGG pathway analysis of the identified proteins (**Table S1**) and metabolites (**Table S2**). Pathway data are in **Table S3**. (**B**) Example showing proteins and metabolites enriched in the arginine-proline pathway. (**C**) Unsupervised (blinded) hierarchical cluster–heat map analysis of the detected protein and metabolite quantities revealing reproducible, systematic, and significant differences between the cell types. Top 40 most significantly dysregulated molecules shown (*p* < 0.05, see quantitative statistics in **Table S3**).

Finally, we applied the technology to ask whether the pluripotent cells harbored detectable differences in proteo-metabolomic activity at such an early stage of development. Although embryonic cells have reproducible tissue fates in the cleavage-stage embryo of *Xenopus*, unbiased and parallel characterization of their proteome and metabolome has not been possible due to a lack of sufficiently sensitive and specific molecular tools. We relatively quantified ~450 proteins and ~80 metabolites between the L-D1 and L-V1 cells in each embryo (N = 4) based on the subcellular HRMS data. Quantification was based on signal abundances in the HRMS datasets, which offer an effective and robust proxy for concentration in CE-ESI-HRMS.^8, 14^ Quantified protein and metabolite signals are presented in **Table S3**. As highlighted earlier, the phenotypes of the cells or the identifier of each embryo were hidden during data analysis to afford bias-free data analysis. **Figure 3C** presents the unsupervised hierarchical cluster analysis (HCA) and intensity heat map of the data based on the top 40 statistically most significantly dysregulated molecules, including 13 metabolites and 27 proteins. Pathway data are listed in **Table S2**. The top dendrogram clusters the samples into two categories in the HCA plot. Upon revealing the identity of the samples, we learned that the groups corresponded to the D1 and V1 cell types. Further, the heatmap revealed notable quantitative molecular differences between the cell types. The left dendrogram organized the proteins and metabolites into two groups in the HCA plot: 20 molecules, containing 10 metabolites and 10 proteins, were more abundant in the L-D1, whereas 20 molecules, encompassing 3 metabolites and 17 proteins were enriched in the L-V1. **Table S3** tabulates proteins and metabolites with statistically (P < 0.05) and biologically (|fold change| ≥ 1.5) significant dysregulation. These intriguing molecular differences would have been lost due to averaging across a large number of cells pooled for classical HRMS approaches.

These previously unknown proteo-metabolomic differences afford new insights into the establishment of cell heterogeneity in the embryo; they also challenge our current understanding of the underlying molecular processes. Differential production of these molecules reveals that asymmetry along the dorsal-ventral, one of the three primary axes of the vertebrate body plan, is set up rather early during embryonic development, when transcriptional heterogeneity is not detectable along this axis based on sequencing of the respective single-cell transcriptomes^36–37^ in *Xenopus*. These findings support our earlier discovery of molecules, such as metabolites^21^, affecting tissue fates via noncanonical mechanisms of action. Although determining the exact biological significance of these findings goes beyond the scope of this study, the data generated in this study may be used to develop hypotheses for experiments targeting specific proteins and metabolites and their functions. Supporting future investigations, our technology is scalable in space and time to other types of cells and different biological models, compatible with complex tissues and live development. It does not escape our attention that our technology can be used to perform multi-‘omics on subcellular organelles. Further, the approach is also adaptable to classical and modern tools of biology and health studies, such as optical microscopy and behavioral assays (as demonstrated in this study) to characterize phenotypes as well as established or contemporary tools of molecular biology, including expression/translation-blocking morpholinos or gene editing by CRISPR-Cas9, to probe function. We anticipate *in vivo* proteo-metabolomic subcellular CE-ESI-HRMS to expand the contemporary toolbox of cell and developmental biology, neuroscience, and health research to understand the molecular basics of the building block of life, the cell, and processes governing the formation of tissues, organs, and entire organisms with complex behavior during state of health and disease.

## Author Contributions

P.N., C.L-B., J.L., and E.P.P. designed the research. C.L-B and R.Q.A. prepared and C.L-B. measured the proteomic samples. J.L., E.P.P., R.M.O, and D.P. prepared and J.L. measured the metabolic samples. J.L., E.P.P., C.S., and P.N. designed and conducted the survival, anatomical, and behavioral assays. P.N., C.L-B., and J.L. analyzed the data, interpreted the results, and wrote the manuscript. All the authors commented on the manuscript.

## Acknowledgments

This research was partially supported by an Arnold and Mabel Beckman Foundation Beckman Young Investigator award (to P.N.), the National Science Foundation under awards DBI-1826932 (to P.N.) and IOS-1832968 CAREER (to P.N.), and the National Institutes of Health under award 1R35GM124755 (to P.N.). The content presented in this work and the conclusions drawn are solely the responsibility of the authors and do not necessarily represent the official views of the sponsors. We are grateful to Dr. Sally Moody (The George Washington University, Washington, DC) for supplying *X. laevis* embryos for the pilot experiments.

