## Supplementary material for "*In Vivo* Subcellular Mass Spectrometry Enables Proteo-Metabolomic Single-cell Systems Biology in a Chordate Embryo Developing to a Normally Behaving Tadpole (*X. laevis*)": SI Document

### MATERIALS AND METHODS

**Materials and Reagents**. All reagents and solvents were obtained at reagent grade or higher purity from Fisher Scientific (Pittsburg, PA) unless otherwise noted. Solutions for MS were prepared using LC-MS grade solvents and reagents (methanol, acetonitrile, water, formic acid, and acetic acid). For CE, bare fused silica capillaries (40/105 µm inner/outer diameter) were purchased from Polymicro Technologies (Phoenix, AZ) and used after initial conditioning (100 mM sodium hydroxide for 5 min, then rinsed with water). CE micro-flow ESI (CE-µESI) for metabolomics employed a leaser-cleaved stainless-steel blunt needle as the spray emitter (130/260 µm inner/outer diameter) from Hamilton Company (Reno, NV). The CE-nanoESI setup for proteomics used a borosilicate capillary (0.75/1 mm inner/outer diameter) tapered on a Blaming/Brown-style capillary puller (P-1000, Sutter Instruments, Novato, CA) and cleaved to a 20-µm tip diameter under a stereomicroscope (model SMZ18, Nikon, Melville, NY).

**Solutions.** The *"metabolite extraction solvent"* was an aqueous mixture of 40% (v/v) acetonitrile (ACN) and 40% (v/v) methanol (MeOH) solution, chilled to 4 °C. The *"protein extraction–digestion buffer"* was 50 mM ammonium bicarbonate in water. For CE, the *"background electrolyte (BGE)"* was 1% (v/v) formic acid for metabolomics and 25% (v/v) MeOH with 1 M formic acid for proteomics. The *"CE-µESI sheath solution"* was aqueous 50% (v/v) methanol containing 0.1% (v/v) formic acid. The *"nanoESI sheath solution"* was 10% (v/v) MeOH containing with 0.05% (v/v) acetic acid.

**Animal Care and Embryology.** All protocols concerning the humane maintenance and handling of vertebrate *X. laevis* animals were approved by the University of Maryland Institutional Animal Care and Use Committee (IACUC No. R-DEC-17-57). Adult male and female frogs were received from Nasco (Fort Atkinson, WI) and maintained in a breeding colony. Embryos were obtained by gonadotropin-induced natural mating of two sets of parents, dejellied in 2% cysteine solution, and cultured in 100 % Steinberg's solution following standard protocols.[1] Two-cell embryos displaying stereotypical pigmentation, size, and location[1] were transferred into a Petri dish coated with 2% agarose containing 100% Steinberg's solution. The embryos were monitored under a stereomicroscope until they reached the 8-cell stage, then the left dorsal-animal (L-D1) and left ventral-animal (L-V1) cells were identified based on reproducible cell-fate maps[2].

**Microprobe Sampling and Sample Processing.** Contents of the L-D1 and L-V1 cells were collected via capillary microsampling[3-6] in randomized order. In this study, the tip of a borosilicate capillary (0.75/1 mm inner/outer diameter tapered to ~20 µm diameter) was fine positioned into the identified cells under control by a three-axis manual micromanipulator (Warner Instruments, Hamden, CT) and guidance by a high-resolution stereomicroscope (SMZ18, Nikon). An ~10 nL portion of the cell was aspirated each time by delivering calibrated pressure pulses (−40 psi) to the microcapillary using a connected microinjector (model PLI-100A, Warner Instruments, Hamden, CT). The subcellular aspirates were expelled into individual microvials and processed differently for metabolomic[5, 7] and proteomic[6] analyses following our established protocols. Metabolites were extracted into 4 µL of *metabolite extraction solvent* (chilled to 4 °C), facilitated by vortex-mixing for 1 min, and the resulting extract was centrifuged to pellet cell debris (8,000 × *g*, 5 min, 4 °C). Proteins were extracted into 5 µL of *protein extraction–digestion buffer*, denatured at 60 °C, and digested to peptides with 0.5 µL of 0.5 µg/µL trypsin (37 °C, 5 h). The resulting digests were dried in vacuum at 60 °C. The resulting metabolite and protein samples were stored at −80 °C until analysis.

**Behavioral Assays.** A portion of the 2-cell embryos was cultured to the tadpole stage to study their visual behavior using a background color preference assay. This assay reproducibly and robustly measures the differential preference of tadpoles to spend time over light and dark backgrounds between Nieuwkoop-Faber Stages 45 and 50.[8] The tadpoles were humanely raised, maintained, and treated following protocols approved by the University of Maryland IACUC (protocol no. R-JUN-20-31). The residence times of each tadpole over the light (white) and dark (black) backgrounds were quantified in a double-tank behavioral system that we built and operated following an established protocol[8]. The movement of each tadpole was recorded on a digital camera (model EOS 70D, Canon USA Inc.) for 2 min on two occasions per day over 2 days. The videos were recorded at 30 frames per second with Full HD resolution (1920 × 1080 pixel$^2$) and using ALL-I compression method. The percentage of time that each tadpole spent in total over the light and dark areas was calculated based on the location of the tadpoles' eyes by manually reviewing the recording, frame by frame, in Windows Media Player version 12 (Microsoft, Redmond, WA). A tadpole was considered to cross the white-black boundary only if both its eyes delineated the boundary.

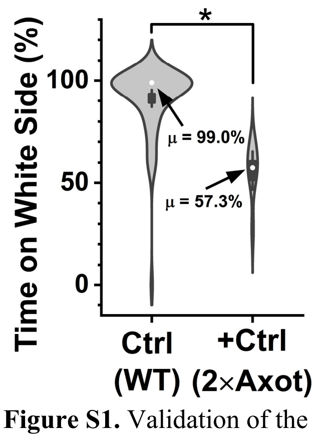

**Figure S1.** Validation of the behavioral assay. Double axotomy of the optic nerves significantly impacted the background color preference of the tadpoles. Key: *$p < 0.05$ (Mann-Whitney test).

Validation. The background color preference assay was validated using a positive control (+Ctrl.) group and by comparing results from our study to those independently established by others.[8] The control group (Ctr.) consisted of wildtype (WT) tadpoles (N = 16). The +Ctr group consisted of WT tadpoles (N = 4) one day after double axotomy of the left and right optic nerves following the protocol established in Ref. [8]. The movement of each tadpole was tracked during the background preference assay, as described earlier. Without visual perception, tadpoles were anticipated to randomly explore the white and black areas, viz. spending ~50% of time over either background color. Indeed, in an independent study, tadpoles post double axotomy spent 49–51% of time over the white background (Day 1–Day 2), whereas the control explored this area in 99%–79% of time with this difference being statistically significant.[8] In our experiments, (**Fig. S1**), the mean time tadpoles spent over the preferred white background was ~56% in the positive control group vs. ~91% in the Ctrl. group with this difference amounting to statistical significance in our measurements (Mann-Whitney test, $p = 1.64\times10^{-5}$). Representative videos are provided for a Ctrl. (**Video S1**) and a +Ctrl. tadpole (**Video S2**). These results validated the sensitivity and robustness of the behavioral assay to support our studies on tadpole behavior.

The behaviors of experimental and control tadpoles were compared. The biological replicate size for the behavioral assay was N = 16 in the Ctr. group (tadpoles raised from nonmicrosampled embryos) and N = 15 in the experimental group (Exp., tadpoles raised from microsampled embryos). **Figure S2** presents representative images of the tadpoles. Power analysis was performed to compute the power to distinguish the wildtype group from the experimental group. Using two sample t-test ($\alpha = 0.05$, sample size = 60), the power was 0.08, which was below the acceptable power required to distinguish the two groups (> 0.8). Together with the Mann-Whitney test, these results indicated that microsampling imposed no detectable impact of statistical significance on the background preference of the tadpoles within the tested sensitivity of the behavioral assay.

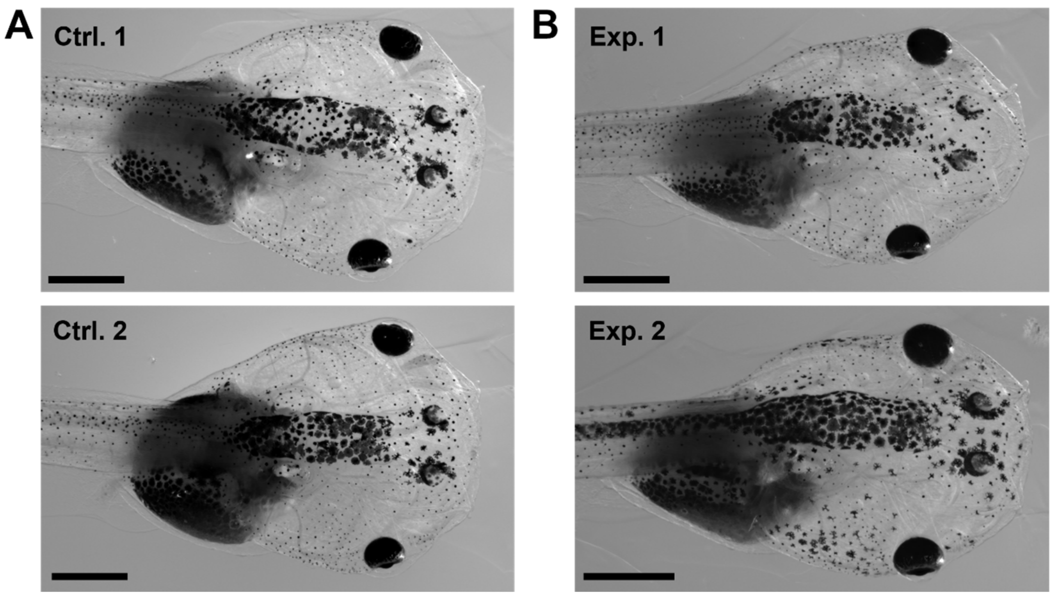


**Figure S2.** Representative high-resolution images of live tadpoles. Close-ups of tadpoles from **(A)** the control (Ctrl.) and **(B)** the experimental (Exp.) group. Scale bars, 1 mm.

**Metabolomic Measurements**. Each subcellular metabolite extract was defrosted, and a 10 nL portion was analyzed on a custom-built CE-ESI platform with a high-resolution time-of-flight tandem mass spectrometer, built and operated according to our established protocols.[5, 9] The instrument was validated for the following analytical figures of merit: lower limit of detection, 60 amol (acetylcholine); linear dynamic range of quantification, 4–5 log order (acetylcholine, methionine, and threonine); reproducibility, <1% relative standard deviation (RSD) with nonlinear time alignment[10] in separation time and <25% RSD in under-the-curve peak area. The experimental settings specific to this study were: CE, 100 cm capillary length and +22.5 kV potential (applied to the capillary inlet); ESI, coaxial sheath-flow CE-ESI interface (blunt-tip metal emitter, 1 µL/min sheath liquid, electrospray at –1,700 V, cone-jet spraying regime); MS, quadrupole orthogonal acceleration time-of-flight mass spectrometer with a collision induced dissociation cell (Impact HD, Bruker Daltonics, Billerica, MS) delivering 40,000 full width at half maximum (FWHM) resolution and <5 mDa m/z accuracy (both single- and tandem stage) using nitrogen gas at 12–18 eV collision energy. Each sample was analyzed in technical duplicate-triplicate.

**Bottom-up Proteomic Analysis by DDA CE-ESI-HRMS.** Each subcellular protein digest was defrosted and individually reconstituted in 2 µL of aqueous 60% acetonitrile containing 0.05% acetic acid. A 10 nL portion of each protein digest was analyzed on a custom-built CE-nanoESI platform equipped with a quadrupole orbitrap high-resolution mass spectrometer. The instrument was built and operated following our established protocols.[11-13] The platform was validated to deliver the following analytical figures of merit: lower limit of detection, 700 zmol (angiotensin II); linear dynamic range of quantification, 4–5 log-order; reproducibility, <10% RSD in migration time and <25% RSD in signal intensity. The experimental settings specific to this study were: CE, 90 cm capillary length and +18 kV potential for 15 min and +15 kV thereafter (applied to the capillary inlet); CE-nanoESI[14], electrokinetic pump supplying 10% MeOH (0.05% AcOH) at +800–1,000 V (applied to the sheath reservoir), cone-jet spraying regime confirmed by a long working distance camera (EO-2018C with Mitutoyo Plan Apo objective, Edmund Optics, Barrington, NJ); and MS, quadrupole orbitrap mass spectrometer (Q-Exactive Plus, Thermo Scientific, Milford, MA).

Experimental conditions of data-dependent acquisition were tailored to fast and efficient separation by CE. Survey spectra were recorded at 35,000 FWHM (*m/z* 200) between *m/z* 350–1,600 (C-trap maximum injection time, 50 ms; AGC target, $1\times10^6$ ions). $MS^2$ scans were triggered for ion signal intensity exceeding $1.5\times10^3$ counts when the charge state was between +2–+7 and exhibited peptide-like isotopic distribution. An exclusion list of ~250 vitellogenin (Vtg) peptides was created based on identifications from a whole embryo "pre-run" to help minimize/avoid the sequencing of abundant yolk-related peptide signals. The $MS^1$ exclusion mass tolerance was 3 ppm. The $MS^2$ scans parameters were as follows: mass resolution, 17,500 FHWM (*m/z* 200); C-trap maximum injection time, 60 ms with AGC target, $5\times10^4$ counts; dynamic exclusion time and mass tolerance, 9 s and 5 ppm; peptide isolation window, 1 Da; collision parameters, HCD in nitrogen at 28% normalized collision energy (NCE). Each protein digest was measured in technical duplicate-triplicate.

To increase identification numbers using the match between run feature, we also measured cell samples using our CE-nanoESI coupled to a Fusion Lumos quadrupole-linear ion trap-orbitrap tribrid mass spectrometer (Thermo Scientific). $MS^1$ parameters were: mass analyzer, orbitrap; mass range, *m/z* 350–1,600; mass resolution; 50,000 FWHM (*m/z* 200); C-trap maximum injection time, 86 ms; automatic gain control (AGC) target, $1\times10^6$ ions. $MS^2$ scans were triggered for ion signal intensity exceeding $1.5\times10^3$ counts, with charge state between 2–7, and exhibited peptide-like isotopic distribution. The $MS^2$ scans parameters were as follows: mass analyzer, ion trap; AGC target, $5\times10^4$ counts; dynamic exclusion time and mass tolerance, 9 s and 5 ppm; peptide isolation window, 1 Da; collision parameters, HCD in nitrogen at 32% normalized collision energy.

**Identification and Quantification of Metabolites and Proteins.** *Metabolites* were identified and relatively quantified following our established protocols.[3, 5, 9] Molecular features (signals with unique *m/z* vs. migration time values) with $S/N \geq 3$ were surveyed between *m/z* 50–500 using a semi-automated approach. For each molecular feature, the under-the-curve peak area was integrated, providing a proxy for concentration. These quantitative metadata were imported to MetaboAnalyst version 5.0[15] for mean-normalization and log-transformation, then exported for integration with the proteomics metadata, before importing of the unified metabolomic-proteomic metadata for statistical data analysis. Select molecular features were identified by matching the accurate mass, isotope distribution, CID fingerprint, and migration time of the unknown signal to data available in metabolomics MS-MS/MS databases (Metlin[16]) or experimentally determined by us in this study or previously[3, 17].

*Proteins* were identified and quantified following established protocols. The MS-MS/MS data were matched against the *X. laevis* proteome using MaxQuant[18] version 1.6 running the Andromeda search engine.[19] The *X. laevis* proteome database was custom built by concatenating the mRNA-derived PHROG1r0 database (downloaded from reference[20]) and the *X. laevis* SwissProt proteome database (downloaded from UniProt on Oct. 2019). The following search parameters were used: no fixed modification; variable modifications, methionine oxidation, and asparagine and glutamine deamidation; minimum peptide length, 5 amino acids; $MS^1$ mass deviation, 5 ppm; $MS^2$ mass tolerance, 20 ppm for first search and 10 ppm for de novo tolerance; search for common contaminants enabled; match between run enabled with 5 min time shift tolerance. A minimum of one unique peptide was required for successful protein identification. Identified common contaminants were manually excluded from proteins identified or quantified in this study. Peptide and protein identifications were filtered to <1% false discovery rate (FDR) calculated against a reversed-sequence decoy database. Proteins were quantified using the label-free quantification (LFQ) approach in MaxQuant using unique and razor peptides.[21] Only proteins with less than 50% missing LFQ values across all the samples were considered as successfully quantified and used for statistical and pathway analysis. Protein LFQ values were mean-normalized and log-transformed in MetaboAnalyst version 5.0[15], then exported for integration with the metabolomics data, before importing of the unified metadata for statistical analysis.

**Experimental Design and Statistics.** For each portion of this study, the number of necessary technical and biological replicates for statistical significance were determined based on a pilot study and the analytical performance metrics of the custom-built CE-ESI-MS platform. Statistical, multivariate, and joint pathway analyses were performed in MetaboAnalyst 5.0. Statistical significance was marked at $p \leq 0.05$ (paired student t-test for normally distributed data). Power analysis (two-sample t-test, alpha value 0.05, hypothetical sample size N = 60) , Student's t-test (parametric, normally distributed data), Mann-Whiney test (nonparametric, non-paired data), and Wilcoxon signed rank test (nonparametric, paired data) were performed in OriginPro 2020b (OriginLab Corp., Northampton, MA). For pathway enrichment analysis, gene and metabolite centered pathway was chosen. Enrichment was measured using hypergeometric test, topology was calculated by degree of centrality, and the integration method used was combined *p*-value (overall).
